# Cross-platform proteomics to advance genetic prioritisation strategies

**DOI:** 10.1101/2021.03.18.435919

**Authors:** Maik Pietzner, Eleanor Wheeler, Julia Carrasco-Zanini, Nicola D. Kerrison, Erin Oerton, Mine Koprulu, Jian’an Luan, Aroon D. Hingorani, Steve A. Williams, Nicholas J. Wareham, Claudia Langenberg

## Abstract

Discovery of protein quantitative trait loci (pQTLs) has been enabled by affinity-based proteomic techniques and is increasingly used to guide genetically informed drug target evaluation. Large-scale proteomic data are now being created, but systematic, bidirectional assessment of platform differences is lacking, restricting clinical translation. We compared genetic, technical, and phenotypic determinants of 871 protein targets measured using both aptamer-(SomaScan® Platform v4) and antibody-based (Olink) assays in up to 10,708 individuals. Correlations coefficients for overlapping protein targets varied widely (median 0.38, IQR: 0.08-0.64). We found that 64% of pQTLs were shared across both platforms among all identified 608 *cis*- and 1,315 *trans*-pQTLs with sufficient power for replication, but with correlations of effect estimates being lower than previously reported (*cis*: 0.41, *trans*: 0.34). We identified technical, protein, and variant characteristics that contributed significantly to platform differences and found contradicting phenotypic associations attributable to those. We demonstrate how integrating phenomic and gene expression data improves genetic prioritisation strategies, including platform-specific pQTLs.

## INTRODUCTION

Proteins are the essential functional units of human metabolism that translate genomic information and enable growth, development, and homeostasis. Naturally occurring sequence variation in the human genome, either in close physical proximity to the protein-encoding gene (*cis*) or anywhere else in the genome (*trans*), has wide-ranging effects on proteins, including, but not limited to, expression, structure, or function, with possible major health consequences^1,2^. Early studies have started to describe the genetic architecture of protein targets measured in plasma but have been small scale or restricted to one platform^3–8^.

Modulating protein abundances or function represents the most common mode of action of drugs^9^ and major pharmaceutical companies now integrate protein quantitative trait loci (pQTLs) into their strategies to identify new drug targets or repurpose existing drugs^10–12^. This has only been possible through the commercial development and application of scalable affinity-based proteomic techniques that can measure thousands of protein targets simultaneously. Projects are now underway to apply these techniques to large-scale studies, such as the UK Biobank^13,14^, which will provide major scientific opportunities. However, information about the consistency of protein measures and pQTLs across platforms is needed to inform the generalisability of genetic findings and strategies for future data integration and meta-analytical approaches.

The deep coverage of the plasma proteome, including thousands of proteins, is possible using large-scale libraries of affinity reagents, with the SomaScan® assay (aptamer-based) and the Olink proximity extension assay (PEA, antibody-based) providing the broadest coverage. Briefly, the SomaScan assay utilizes short single-stranded oligonucleotides, which are chemically modified to increase affinity to specific protein targets and DNA microarray technology is used to quantify the number of aptamers bound to protein targets^15^. Olink relies on monoclonal or polyclonal antibodies labelled with single oligonucleotides that create pairs of antibodies binding to different epitopes of the protein target. The oligonucleotides hybridise only if a matched pair of antibodies bind, and the resulting short DNA fragment is measured using qPCR or next generation sequencing^16^. Despite measurement units, i.e., relative intensities, not being directly comparable between platforms, analysis of correlations and rank based variation between platform is scale-free and, further, orthogonal evidence from genetic variation at protein-encoding genes (*cis*-pQTLs) can be used to compare platforms.

Both techniques rely on conserved binding regions of the protein target, epitopes, to provide reliable estimates of protein concentrations and protein altering variants mapping to epitopes haven been widely recognized to possibly introduce binding artefacts^1,17,18^. While we and others^3,5,18,19^ have recently demonstrated that pQTLs can successfully be replicated across platforms for a selected set of overlapping proteins, this has not been systematically evaluated across hundreds of protein targets than can now be mapped across the latest versions of these platforms.

Here we assess 871 proteins targeted by both the Somalogic and Olink platforms and measured in up to 10,708 individuals, including overlapping measurements by both technologies in a subset of 485 participants. We use a machine learning approach to identify technical parameters and protein characteristics that contribute to variation between platforms. We identify hundreds of pQTLs and systematically assess their consistency in a reciprocal design, that is, by taking each pQTL forward for assessment irrespective of the discovery platform, generating a unique benchmark for future studies.

## RESULTS

We used the SomaScan v4 platform (SomaLogic Inc., Boulder, Colorado, US) to measure protein abundances of 4,775 unique human protein targets (covered by 4,979 unique aptamers) from frozen EDTA-plasma samples in 12,345 participants in the Fenland study. We assessed 1,069 protein targets based on 1,104 measures across 12 Olink® Target 96-plex panels, based on the proximity extension assay (PEA) technology using the same EDTA-plasma samples from 485 Fenland study participants. Measurements were performed by the manufacturers and methods have previously been described in detail^19,20^ and are provided in the Methods section.

We identified overlapping protein targets between both techniques using either UniProt identifiers (www.uniprot.org) or based on the same encoding gene as provided by the manufacturers. Where multiple measurements were available for a protein assayed on multiple Olink panels, we selected one of the protein measures from one of the panels at random for two reasons. Firstly, Olink uses the same type of antibodies irrespective of the panel and secondly, the average correlation was 0.90 (range 0.68-0.99) for the same protein target across different panels. We kept each SOMAmer reagent matching to one Olink reagent for downstream analysis, since they bind to distinct structural characteristics of the protein target^15^. This procedure yielded 937 unique SOMAmer – Olink measurement pairs, comprising 871 unique protein targets (**Fig. 1 and Supplementary Tab. S1**).

**Figure 1.**
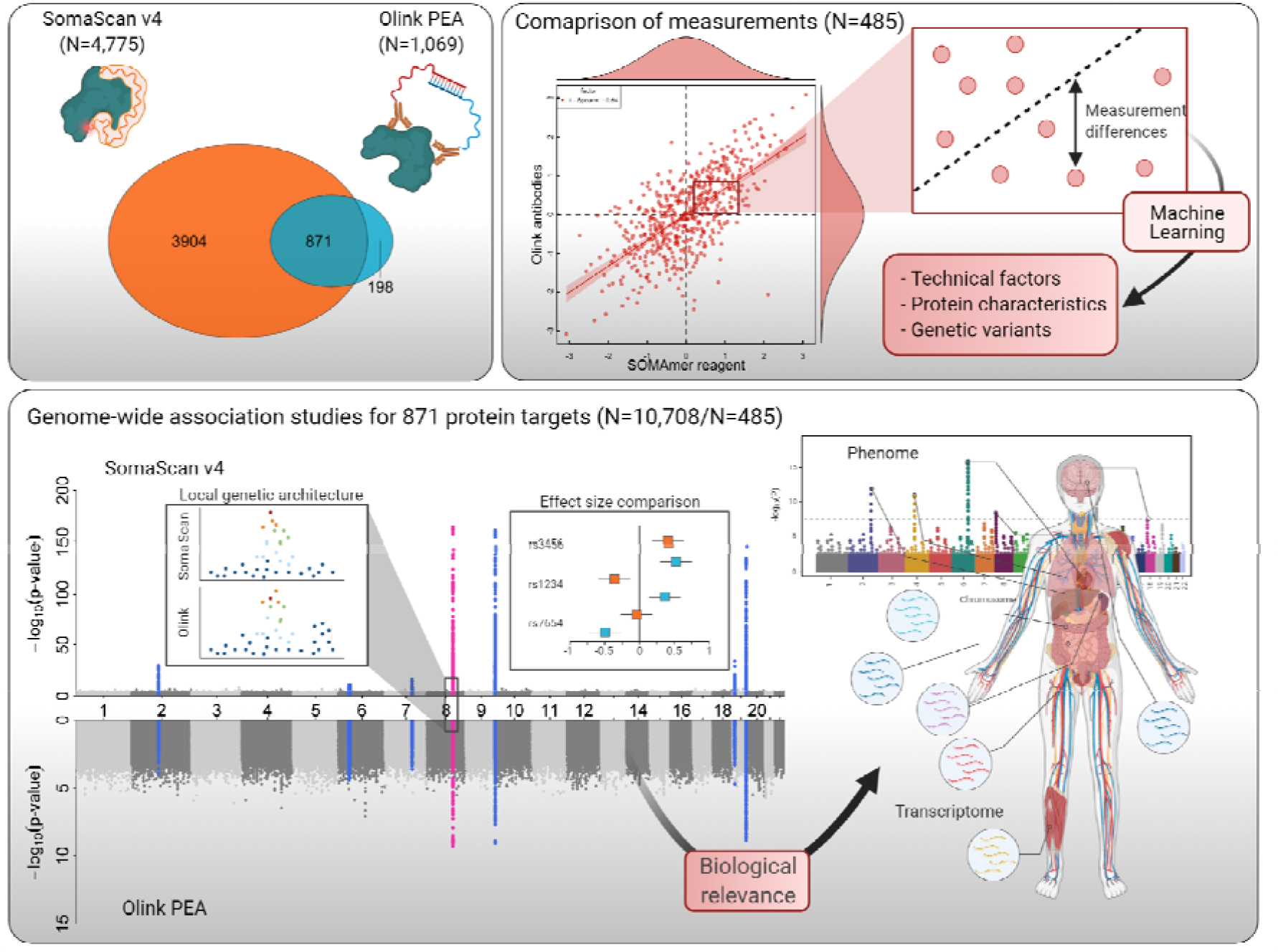
Scheme of the study design. The Venn diagram displays the overlap in protein targets captured by the SomaScan assay and the Olink proximity extension assay (PEA). Modes of binding to the protein target are depicted simplified next to each ellipse. Correlation coefficients were used to compare both technologies and factors possibly accounting for measurement differences and low correlation coefficients examined in a subset of 485 individuals with overlapping measurements. For the set of 871 common protein targets genome-wide association analysis were performed in 10,708 (SomaScan assay) and 485 (Olink PEA) participants of the Fenland cohort. Correspondence of genetic associations was analysed by examining local genetic architecture, comparison of effect estimates, and evaluation of phenotypic consequences.

### Technical factors affecting correlations between protein targets

The median Spearman correlation coefficient between overlapping protein targets was 0.38 (IQR: 0.08-0.64), with large variation (range: −0.61 to 0.96), including examples with high (Leptin, r=0.95), absent (Interleukin-12, r=0.02), and inverse correlations (Heat shock protein beta-1, r=-0.48) **(Fig. 2a and Supplementary Tab. 1)**. We tested for systematic variation among correlation coefficients due to technical factors or protein characteristics using a random forest-based feature selection algorithm (see **Methods**). We identified assay characteristics, including values below the detection limit of the assay, the affinity of the SOMAmer reagent to its protein target (‘apparent Kd’), or the proportion of measurements far off from the median value (‘%-outlier SomaScan/Olink’ – median ±5*MAD), to be more relevant to explain varying correlation coefficients compared to any structural properties of the assayed protein targets (**Fig. 2**). Proteins with a transmembrane domain showed on average lower correlations compared to those without (**Fig. 2)**. We suspect that SOMAmer reagents or antibodies binding to the part of the plasma protein excluding the transmembrane domain may measure both intact and post-translationally modified proteins, those that are generated after proteolytic removal of the ectodomain^21^. Inflammatory mediators, such as tumour necrosis factor alpha, are activated by proteolytic cleavage from the transmembrane domain and hence the ability to specifically target and distinguish this active fraction of the protein target may be relevant to the studying of its pathological relevance. Further, selection of affinity reagents against proteins with transmembrane domains might be complicated by the incomplete *in vitro* folding of synthetic peptides.

**Figure 2.**
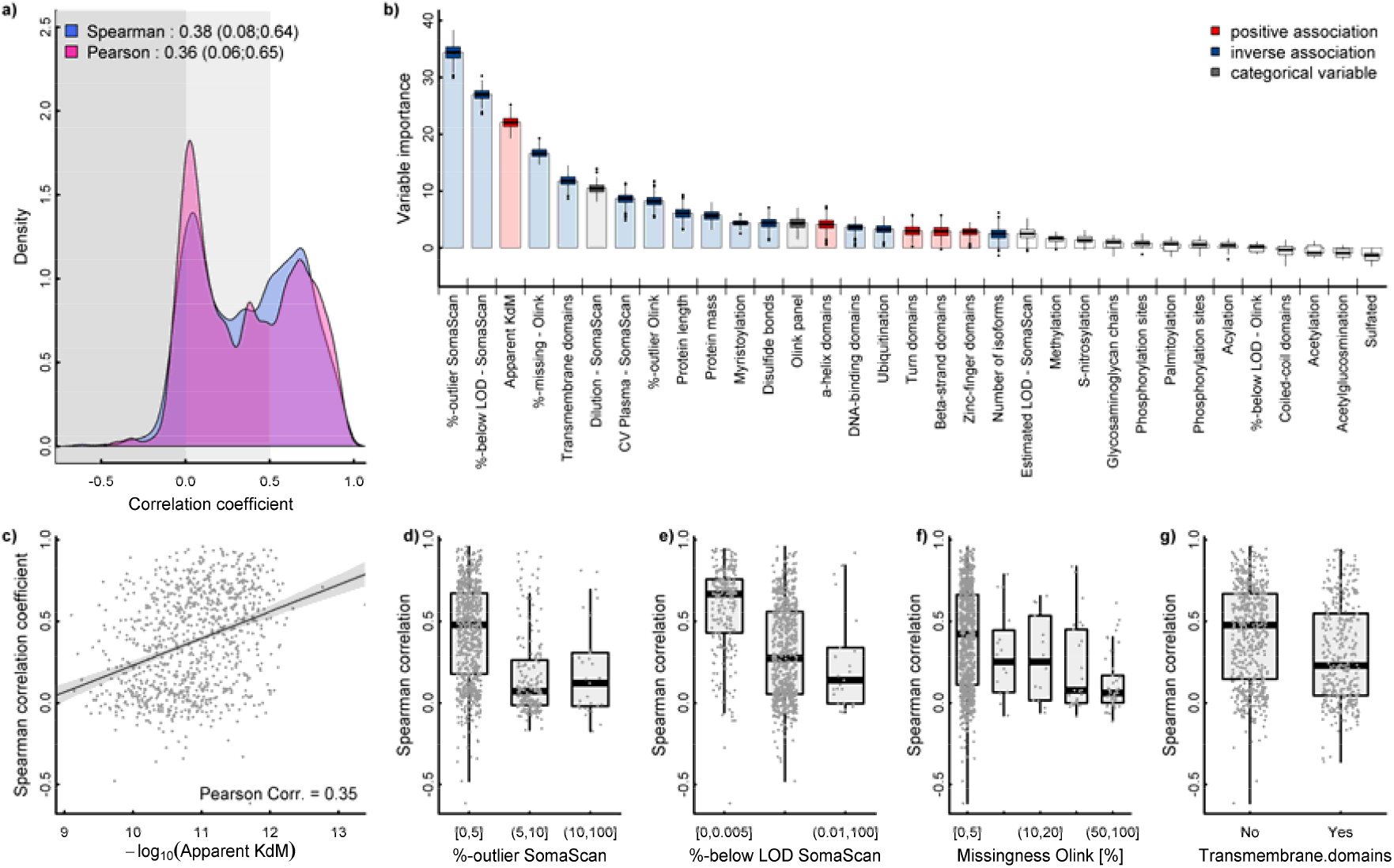
Summary of correlations between measurements on both platforms. **a)** Distribution of correlation coefficients across 937 mapping aptamer – Olink measure pairs (n=871 unique protein targets). **b)** Importance measures derived from a random forest-based variable selection procedure to predict Spearman correlation coefficients including technical factors and protein characteristics. Coloured box plots indicate variables for which the importance measure remained significant after accounting for multiple testing. **c-g)** Individual-level plots of correlation coefficients for the most important characteristics. %-below LOD = Fraction of measurement values below the detection limit of the assay.

We did not find evidence for a systematic pathway bias for either of the two technologies in that protein targets with lower correlations (r<0.2) were not enriched for any particular biological pathway.

### Variation of genetic effect estimates between platforms

To systematically test for cross-platform consistency of protein-quantitative trait loci (pQTLs) we performed a reciprocal comparison of effect estimates of genome-wide association analysis of 871 common protein targets using the SomaScan v4 assay (N=10,708, p<1.004×10^−11^) with 12 Olink panels (N=485, p<4.5×10^−11^, **Fig. 3**) in the Fenland study. This analysis overcomes the biased assessment of previous one-way or within platform replication efforts^4,5,18^. To test the potential influence of sample size on this comparison, we additionally compared the SomaScan-derived pQTLs to published genetic effect estimates for 90 protein targets from the Olink CVD-I panel including up to 22,000 participants from the SCALLOP consortium^8^.

**Figure 3.**
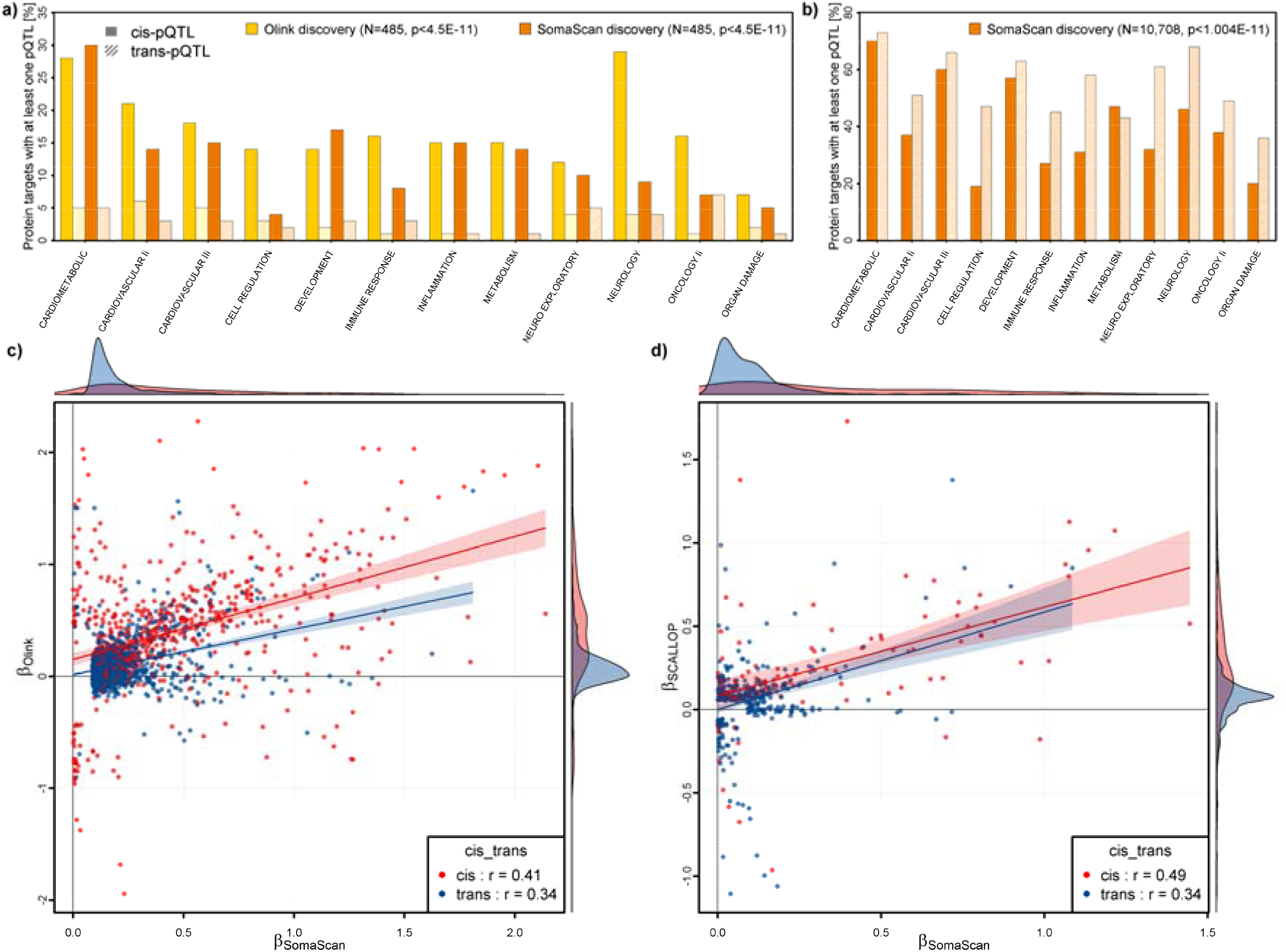
Results from genome-wide association analysis and reciprocal look-up. **a)** Fraction of protein targets with at least one protein-quantitative trait loci (pQTL) in *cis* (opaque) or *trans* (shaded) using either Olink (yellow) or the SomaScan assay (orange) in an overlapping set of 485 participants, grouped by Olink panel. Numbers refer to the set of 871 protein targets measured by both techniques. **b)** The fraction of protein targets with pQTLs in the entire Fenland cohort (N=10,708) based on the SomaScan assay. **c)** Comparison of beta estimates from linear regression models across 816 corresponding SOMAmer - Olink pairs (n=770 unique protein targets) with at least one genome-wide associated genetic variant for either of the two, including 1,267 distinct genetic variants (R^2^<0.8). Colouring is based on the genomic location of genetic variants. Red indicates variants close to the protein encoding gene (*cis*, ±500kb) and blue otherwise. **d)** Comparison of beta estimates from linear regression models across 85 corresponding SOMAmer - Olink pairs (n=77 unique protein targets) with at least one genome-wide associated genetic variant for either of the two, including 428 distinct genetic variants (R^2^<0.8). Genetic variants for Olink measures were derived from the most recent SCALLOP effort covering the CVD-I panel^8^.

We identified a total of 1,923 SOMAmer - Olink - genetic variant triplets (N=608 in *cis*, N=1,315 in *trans*, **Supplementary Tab. 2**) with evidence from either platform, including 816 SOMAmer reagents, 770 Olink measures, and 1,267 single nucleotide variants (SNVs), following pruning of variants in high linkage disequilibrium (LD, R^2^>0.8). The correlation of effect estimates was higher for *cis*-pQTLs (r=0.41) than *trans*-pQTLs (r=0.34) and was considerably lower than those reported in previous studies which did not perform bidirectional assessment or inflated correlation estimates by not aligning effect estimates to either the protein-increasing or −decreasing allele and thereby ‘artificially’ increasing the scale of observed effect estimates^5,19^ (**Fig. 3**). Correlations of genetic effect estimates differed according to observational correlations between targets, with high correlations (0.68 and 0.75 for *cis*- and *trans*-pQTLs, respectively) seen across the 351 protein targets with observational correlations r≥0.5 (**Supplementary Fig. 1**), but no correlation (r=-0.07 for *cis*-pQTLs and r=-0.10 for *trans*-pQTLs) for those 265 with low observational correlations (r<0.2).

Results were consistent (r=0.49 for *cis*-pQTLs and r=0.34 for *trans*-pQTLs, **Fig. 3**) when using summary statistics from up to 22,000 participants for the subset of Olink CVD-I panel proteins (**Supplementary Tab. 3**) and comparing 496 SOMAmer - Olink - SNV triplets (N=95 in *cis*, N=401 in *trans*).

### Cross-platform pQTLs are target-dependent

We collapsed pQTLs discovered by either platform using a distance-based threshold (±500kB, **Fig. 4**) to define shared (‘cross-platform’) versus ‘platform-specific’ pQTLs. This procedure resulted in 479 (N=333 in *cis*, N=146 *trans*, 390 protein targets, **Supplementary Tab. 4**) genomic region – protein target combinations for which we had sufficient statistical power to replicate effects, that is, pQTLs observed in the larger SomaScan study that had at least a p-value<10^−5^ when restricting the analysis to the sample of 485 participants with overlapping measurements (see **Methods**).

**Figure 4.**
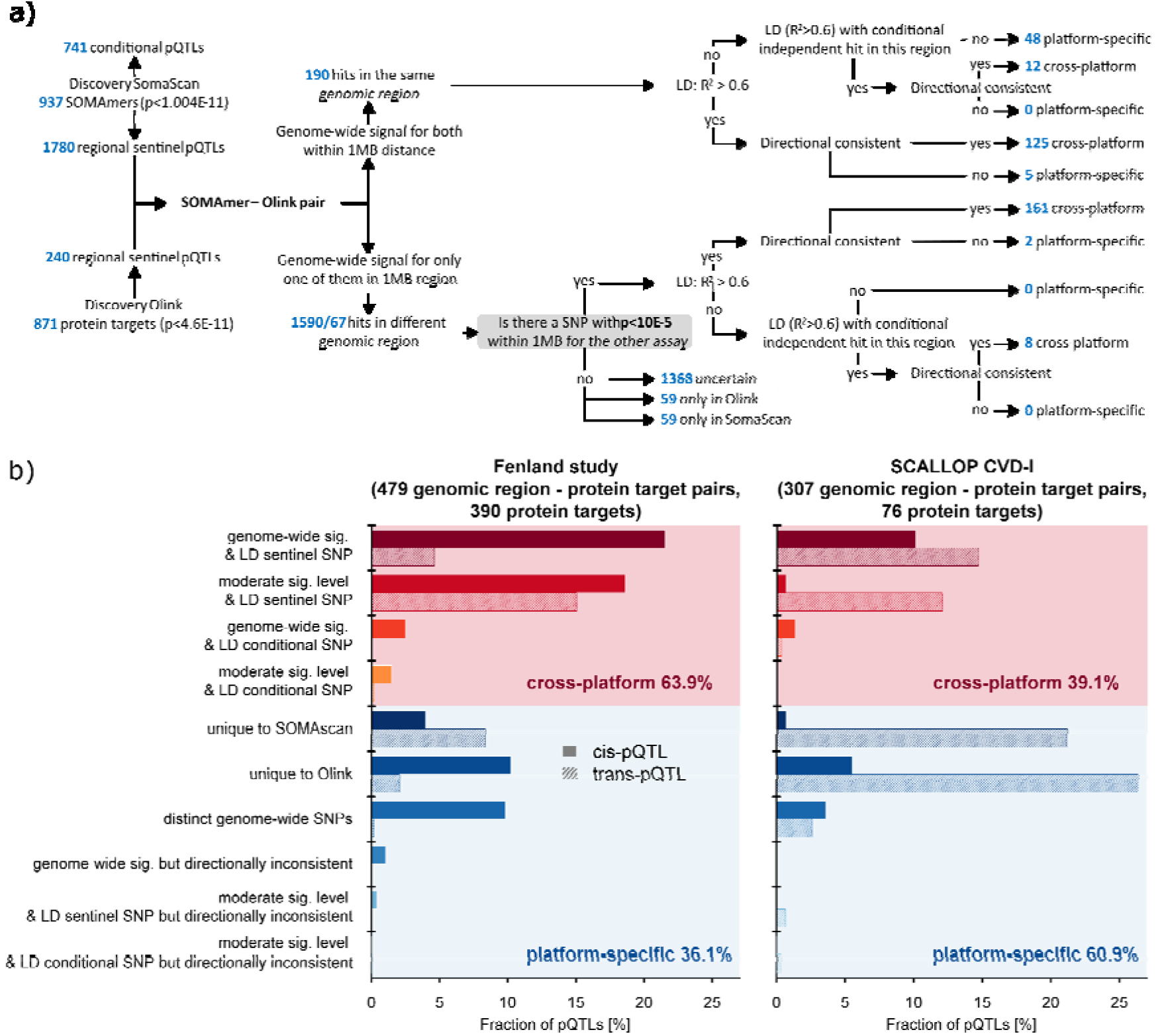
Cross-platform agreement of genomic region – protein target associations. **a** Workflow to determine shared (‘cross-platform’) and platform-specific effects of protein-quantitative trait loci (pQTLs) between SomaScan and Olink based in the Fenland study. **b** Summary of platform agreement for 479 genomic region – protein target associations with sufficient power among the Fenland subsample with available Olink measures (left plot) and 307 genomic region – protein target associations with sufficient power in the Fenland SomaScan study and the SCALLOP CVD-I consortium (right plot).

We applied the following criteria to consider a pQTL/genomic region to be shared across both platforms: 1) genome-wide significance in either discovery approach of the same SNV or a proxy in high LD (R^2^>0.6) and/or sufficient effect strength to be detected in the smaller Olink sample, and 2) to be directionally concordant (**Fig. 4**). We further performed a regional look-up (±500kB) if the regional sentinels for the SomaScan assay and Olink were not in LD with the respective lead variant and tested if a conditionally independent pQTL in the same region may align (**Fig. 4**). We identified 306 (63.9%) cross-platform genomic region - protein target associations with approximately similar fractions for *cis* and *trans*-pQTLs (**Fig. 4**). Among those were 7 regions for which two independent pQTLs (R^2^<0.1) were shared between SomaScan and Olink, but with different ranking in effect strengths, and further 13 regions for which out of two SomaScan signals only the secondary signal at the locus was also seen with Olink. The remaining 36.1% genomic region - protein target associations were platform-specific because they were either 1) only evident for one of the two assays (24.6%, N=59 for the SomaScan assay and N=59 for Olink), or 2) showed evidence for distinct genetic signals at the same locus (10%, 48 pairs).

We identified seven pairs for which the lead pQTL was shared but with opposite directions of effect for the same protein target or its isoforms (**Supplementary Fig. 2 and Supplementary Tab. 4**). For instance, the missense variant rs1859788 (p.G78R, AF=31.7% for the A-allele) in *PILRA* was the lead *cis*-pQTL for Paired immunoglobulin-like type 2 receptor alpha (PILRA) for the Olink measure (beta=-0.74, p<3.48×10^−29^) and we found positive associations with two SOMAmer reagents targeting soluble isoforms of the same protein (6402-8 targeting isoform FDF03-deltaTM (beta=1.26, p<2.67×10^−5193^) and 10816-150 targeting isoform FDF03-M14 (beta=1.26, p<1.53×10^−5360^), but not the SOMAmer reagent designed to target the extra-cellular domain of the canonical protein (8825-4, beta=0.004, p=0.75). Using statistical colocalisation we provide strong evidence of a genetic signal shared between all three different protein measures and Alzheimer’s disease (**Supplementary Tab. 5 and Supplementary Fig. 3**), in line with the A-allele of rs1859788 having been identified as protective for Alzheimer’s disease. PILRA is an inhibitory receptor expressed in dendritic and myeloid cells^22^ and p.G78R was shown to reduce signalling *via* reduced ligand binding, likely modulating microglia migration and activation in the brain^23^. G78R is located in the extracellular-domain common to all three forms of PILRA^22^. Therefore, the positive effect directions of the SOMAmer reagents targeting the two isoforms in the absence of an association with the canonical protein suggest aptamer binding affinity introduced by p.G78R being restricted to the soluble isoform. However, our results cannot distinguish which isoform the polyclonal Olink antibodies target and whether the inverse association reflects reduced binding affinity to the variant protein of at least some of them. We identified similar examples with possible downstream consequences for phenotypic interpretation, including Hepatoma-derived growth factor and HDL-cholesterol levels or Intracellular adhesion molecule 1 and lymphocyte cell count (**Supplementary Tab. 5**).

To test the influence of an unbalanced design, we performed a sensitivity analysis including 307 genomic region - protein targets pairs (N=67 *cis*, N=240 *trans*, N=76 protein targets, **Supplementary Tab. 5**) overlapping with the SCALLOP CVD-I panel GWAS summary statistics. We identified 120 (39.1%) of the pairs as cross-platform, with higher rates in *cis* (55.2%) compared to trans (24.5%) (**Fig. 4**). The higher fraction of platform-specific pairs in *trans* (157 out of 187, 83.9%) might be best explained by two factors. Firstly, variants in *trans* might increase DNA-binding affinity of abundant circulating proteins such as complement factor H (rs1061170 within *CFH*) or Butyrylcholinesterase (rs1803274 within *BCHE*) possibly interfering with SOMAmer reagents^19^, and, secondly, reflect study-specific handling of blood samples like rs3443671 within *NLRP12*, which might only be identified as a pQTL as a result of white blood cell lysis. Out of the 140 platform-specific *trans*-pQTLs, 26 and 25, respectively, were likely attributable to those reasons.

### Identification of factors for cross-platform pQTLs

To identify factors that are associated with pQTLs that are shared across platforms as opposed to those that are platform-specific, we used logistic regression models to systematically test the odds of platform-specificity for 22 factors, including functional annotation of variants, associations with diverse phenotypic traits, gene expression QTL (eQTL), and protein characteristics. We considered three control groups: 1) protein targets with distinct pQTLs in the same genomic region, 2) pQTLs unique to the SomaScan assay, and 3) pQTLs unique to the Olink assay (**Fig. 5 and Supplementary Fig. 4 and Supplementary Tab. 7-9**). In general, the likelihood of a pQTL being shared across platforms was greater compared to either of the three control groups and in both the Fenland and SCALLOP data when the correlation between measurements (‘observational correlation’) and the binding affinity of the SOMAmer reagent was higher (**Fig. 5 and Supplementary Fig. 4**).

**Figure 5.**
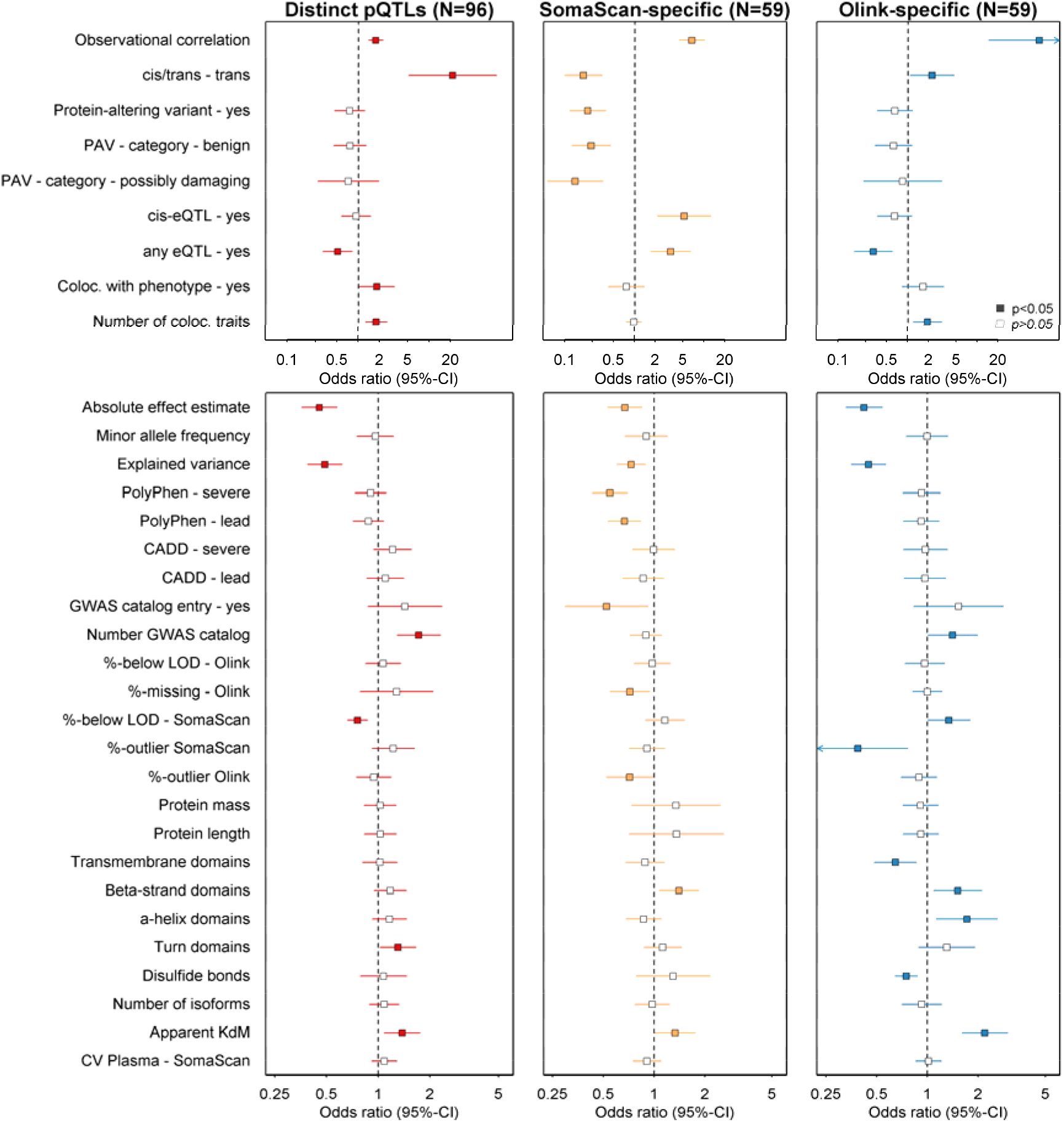
Factors associated with pQTLs that are shared across platforms compared to three sets of platform-specific controls. Odds ratios and 95%-confidence intervals for factors associated with cross-platform protein quantitative trait loci (pQTL) across the SomaScan v4 and Olink assays. The panels are based on 540 variant – protein target pairs (306 shared, 234 platform-specific) with sufficient power for replication in the Fenland sample. PAV = protein altering variant; eQTL = expression quantitative trait loci; Coloc. = colocalisation; GWAS = genome-wide association analysis; LOD = limit of detection; KdM = estimated apparent dissociation constant (Kd) of SOMAmer reagents in molar units (M)

We identified a few factors that were significantly associated only when using specific control groups. For example, LD with a (benign) PAV decreased the odds for cross-platform pQTLs only when comparing to SomaScan-specific pQTLs, which might be best explained by the putative change in shape of the protein target among carriers of the alternative allele of PAVs and the reliance of SOMAmer reagents on a conserved shape of the protein target (**Supplementary Tab. 10**). Strong effects of single genetic variants on assay results, indicated by the factor “%-outlier SomaScan”, may even mask associations that would otherwise be expected from the protein target, that is only seen with the Olink assay. Further, colocalization with (cis)-eQTLs and phenotypic traits was associated with a higher likelihood of cross-platform pQTLs comparing to SomaScan-specific and distinct pQTLs, respectively (**Fig. 5**).

Platform-specific pQTLs with strong evidence for colocalization with a phenotypic trait may provide evidence about the biological relevance of the pQTL. Therefore, exploring those associations may provide insights that would otherwise be hidden if only one platform was analysed. Out of 17 and 47 *cis*-pQTLs unique to the SomaScan and the Olink assay, respectively, in the Fenland study, three and 14 had either been reported in the GWAS catalog or colocalised in a phenome-wide scan (Supplementary Tab. 7-8). While we cannot completely rule out that *cis*-pQTLs attributed solely to the SomaScan assay might not have been observed with Olink due to limited samples size, the 14 *cis*-pQTLs unique to Olink which are demonstrable in only 485 samples suggest that these are strong phenotypic links. For instance, the *cis*-pQTLs rs11589479 (ADAM15) and rs34687326 (SLAMF8) colocalised (posterior probability for a shared genetic signal >90%) with Crohn’s disease (**Supplementary Tab. 5**), of which the missense variant rs34687326 (p.Gly99Ser) within *SLAMF8* has been identified as a risk factor for inflammatory bowel disease as well^24^. We observed a similar distribution of unique *cis*-pQTLs in the larger SCALLOP comparison, with two and 15 *cis*-pQTLs unique to the SomaScan assay and SCALLOP (Olink), respectively, of which one (rs140934622 for IL-27 with the SomaScan assay, in LD, R^2^=0.96, with a lead signal for intelligence^25^) and seven (e.g., rs4512994 for sRANKL on Olink, which is a known susceptibility locus for bone mineral density^26^) had a link to phenotypic traits.

IL-27 is an inflammatory protein and encoded at the two distinct genes *IL27* (IL-27p28 cytokine subunit) and *EBI3* (soluble EBI3 essential for correct folding and secretion)^27^. We identify a SomaScan-specific variant located at *IL27* (rs140934622 on 16p11) and an Olink-specific variant at *EBI3* (rs4905 on 19p13), both of which were in strong LD (R^2^>0.99) with missense variants (rs181206 - p.L119P and rs4740 - p.V201I). It is possible that both missense variants might: 1) differentially affect heterodimerization of the two different gene products required to build IL-27 or 2) both assays have a critical binding epitope on the respective subunit and binding of the affinity reagent is impaired by PAVs.

We considered the identification of secondary signals conditioning on the lead *cis*-pQTL as a strategy to overcome strong platform-specific pQTLs motivated by a subset of 12 protein targets for which not the lead but the secondary *cis*-pQTL was shared across platforms. For three out of those targets, this approach let to the observation of additional phenotypic associations, including IGFBP-3, for which a *cis*-pQTL colocalised (PP>92%) with systolic blood pressure, or FBLN3, which colocalised (PP>80%) with 24 anthropometric traits and operation codes related to hernia (**Supplementary Tab. S5**). Doing the same across 36 protein targets with *cis*-pQTLs unique to SomaScan and evidence for a secondary signal (p<5×10^−8^) revealed three protein targets with a high posterior probability of a shared genetic signal with phenotypes (PP>80%), including CD58 and primary biliary cirrhosis (**Supplementary Tab. S5**).

### pQTLs account for measurement differences

We identified 22 instances in which the correlation coefficient between measurements of the same protein target across both platforms significantly differed by genotype (false discovery rate<20% for an interaction term), including pQTLs in *cis* and *trans* (**Fig. 6**). That is, once carriers of the minor or major allele were excluded, the correlation coefficient between both assays improved. For example, stratifying observational correlations by the two weakly related (R^2^=0.23, D’=0.93) lead *cis*-pQTLs for YKL-40 (rs2071579 - SomaScan, rs4950928 - Olink) raised the correlation coefficients across all categories up to 0.95 (range: 0.56 - 0.95) for carriers of the minor C allele (MAF=45.7%) of rs2071579 (**Fig. 6**). Rs2071579 is in LD with a possibly damaging missense variant (rs88063, R^2^=0.99, CADD score = 22.9, p.R145G), located at a predicted protein-protein interaction site in a highly conserved region of the protein. However, none of the pQTLs has been genetically linked to phenotypes other than the protein itself.

**Figure 6.**
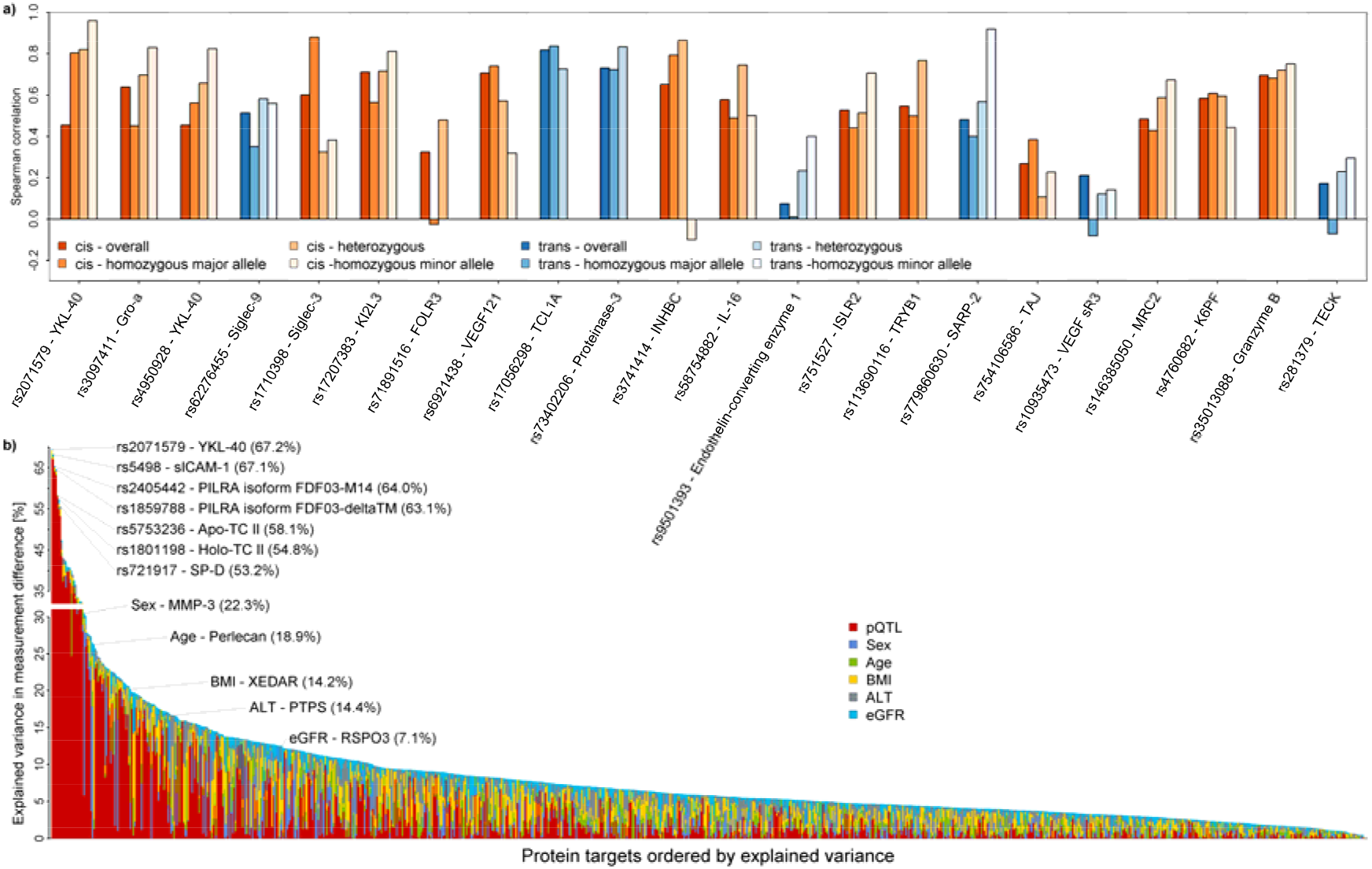
Genetic variants modulate correlation coefficients and explain measurement differences between both assays. **a)** Spearman correlation coefficients stratified by genotype. The first bar in each column indicates the overall correlation, and the three successive bars indicate the correlation among homozygous carriers of the major allele, heterozygous carriers, and homozygous carriers of the minor allele (if any). Colours indicate whether the pQTL was in *cis* (orange) or *trans* (blue). Protein target – pQTL pairs were selected based on a linear regression model (see Main text). *b)* Protein targets ordered by the amount of variance explained in the differences between measurements. Contribution of protein quantitative trait loci (pQTL), age, sex, body mass index (BMI), plasma alanine aminotransferase activities (ALT), and estimated glomerular filtration rate (eGFR) are given in colours. Selected protein targets are annotated.

While such findings in *cis* are likely to indicate possible interference with binding epitopes, variants in *trans* act through various pathways (**Supplementary Tab. 10**). For example, variants mapping to ubiquitously expressed glycosyltransferases may act through alerted glycosylation of protein targets affecting the accessibility for affinity reagents. We observed two such examples, namely rs281379 (associated with TECK and in LD, R^2^=0.83, with a missense variant in *FUT2*) and rs779860630 (associated with SARP-2 and intronic of *ABO*) mapping to genes encoding glycosyltransferases. Another possibility is a higher affinity for RNA or DNA binding of the gene product conferred by the genetic variant, similar to what was discussed previously for variants mapping to *CFH* and *BCHE*. We observed rs9501393 (MAF=13.5%) modulating the correlation coefficient of Endothelin-converting enzyme 1 (**Fig. 6**). Rs9501393 is in strong LD (R^2^=0.94) with a missense variant of uncertain significance in *SKIV2L* (rs449643, p.A1071V) encoding an RNA helicase, a protein with high affinity to bind to RNA or single-stranded DNA oligomers.

We next identified factors that influenced measurement differences at the individual participant data level, considering pQTLs as well as phenotypic measures that could have an impact on protein abundances, namely age, sex, body mass index (BMI), estimated glomerular filtration (eGFR; calculated from serum creatinine, age, and sex), and plasma alanine transaminase activities (ALT). The combination of all factors explained a median amount of 5.6% (IQR: 3.5% - 9.2%) of the differences in measurements reaching values of up to 69.4% for YKL-40 (**Fig. 6 and Supplementary Tab. 11**). For 211 (23%) out of 814 protein targets with at least one pQTL, the pQTL accounted for most of the explained variance (median: 1.0%, IQR: 0.2% - 3.4%), including 85 protein targets with >10%. The strong contribution of certain genetic variants aligns with the results for platform-specific *cis*- and *trans*-pQTLs outlined above.

## DISCUSSION

Identification of DNA sequence variants modulating protein levels or activities and shared with disease loci can identify disease-causing mechanisms and help to prioritize new and repurpose existing drug targets^10^. To inform and advance such strategies, comparison across different measurement techniques can not only validate identified signals but help to better understand the potential biological relevance of platform-specific signals. Broad, systematic assessment of this across many protein targets has until now been hindered by limited overlap across different proteomic platforms. Previous smaller scale studies^3,5,18^ have performed unidirectional validation of pQTLs for a selected set of protein targets and reported inflated correlation estimates due to missing alignment of effect directions to the protein increasing or decreasing allele, thereby introducing an artificially large reference range. We provide the largest systematic identification and characterisation of pQTLs shared across platforms and those that are platform-specific pQTLs based on reciprocal, that is, bidirectional assessment of the two most comprehensive techniques covering 871 overlapping protein targets. We show that the majority of pQTLs are shared across platforms (64%) but with substantially lower correlations than previously reported in *cis* and *trans*. We identify factors associated with platform-specific pQTLs for both platforms, which can directly help to inform strategies for prioritising pQTLs in academic and pharmaceutical efforts that have used either platform at scale, in particular for the thousands of protein targets only assayed by the Somalogic platform and providing unprecedented breadth for discovery studies.

We provide multiple examples for platform-specific pQTLs with strong evidence of a shared disease signal, which, in case of the SomaScan assay, could suggest a biological link *via* the shape of the protein rather than an effect mediated by the abundance of the protein target (**Supplemental Tab. 10**). This has important implications for protein level based casual inference techniques such as Mendelian randomization, where genetic instruments acting in *cis* are used to typically proxy abundance rather than function of the encoded protein, and where such findings can be used for drug target validation, if the encoded protein is druggable. However, strong platform-specific signals can also hide signals that would otherwise be shared across platforms. We demonstrate that the use of conditional association statistics, upon the lead pQTL in the region, provides a strategy to recover relevant biological information.

We identify several characteristics affecting the correlation between both assays, including technical variation, certain protein characteristics, and a strong effect of genetic variants (**Fig. 7**). However, the lack of full technical details of the assays that are not in the public domain as they are commercially sensitive and general methodological differences between the assays did not permit a more rigorous assessment of non-biological factors. This includes the similarity of synthetic peptides used to select bindings reagents or a measure of binding affinity for antibodies, which might likely yield additional insights into possible differences. Incorporation of complementary techniques such as mass spectrometry may help to resolve some of these issues^28^, for example by linking a pQTL to an actually measured peptide sequence, which would provide important scientific opportunities if the approach can be applied at scale. In addition, structural characterization of proteins bound to affinity reagents using mass spectrometry has the potential to identify the concrete protein species bound to the affinity reagent^4,18^.

**Figure 7.**
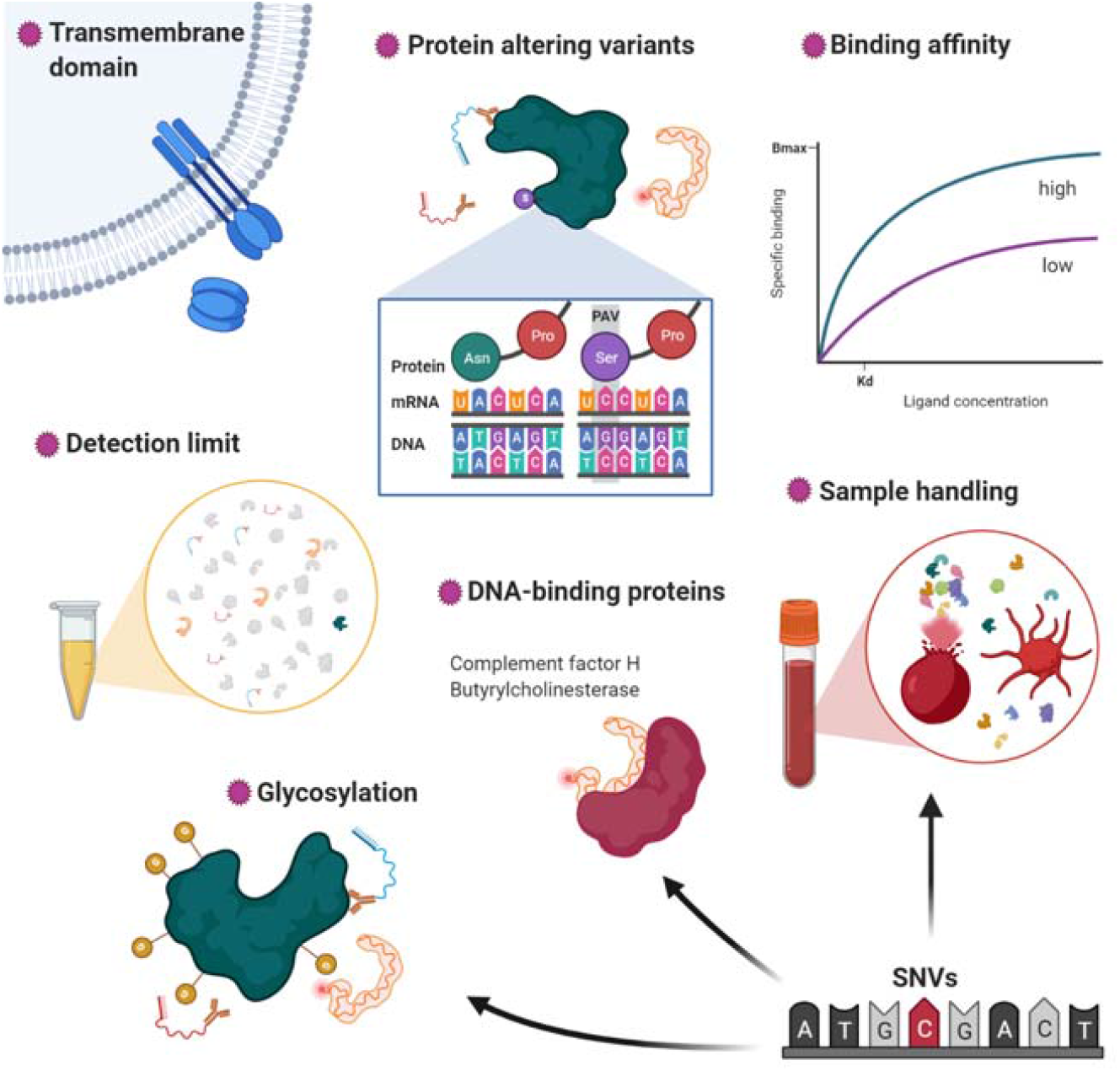
Sources of variation. Graphical summary of factors contributing to variation in the affinity-based discovery of the plasma proteome. PAV = protein altering variant; SNV = single nucleotide variant

## METHODS

### MRC Fenland cohort

The Fenland study is a population-based cohort of 12,435 participants, predominantly of White British ancestry, born between 1950 and 1975 who underwent detailed phenotyping at a baseline visit between 2005 and 2015. Participants were recruited from general practice surgeries in the Cambridgeshire region of the UK. Exclusion criteria were clinically diagnosed diabetes mellitus, inability to walk unaided, terminal illness, clinically diagnosed psychotic disorder, pregnancy, or lactation. The study was approved by the Cambridge Local Research Ethics Committee (NRES Committee – East of England Cambridge Central, ref. 04/Q0108/19) and all participants provided written informed consent. Participants in the study were on average 48.6 years old (standard deviation: 7.5 years) and 53.4% of them were female, as previously described^29^.

### Proteomic measurements

Proteomic profiling of fasting EDTA plasma samples from 12,084 Fenland Study participants collected at the baseline visit was performed by SomaLogic Inc. (Boulder, US) using an aptamer-based technology (SomaScan v4 assay). Relative protein abundances of 4,775 human protein targets were evaluated by 4,979 aptamers (SomaLogic V4) and a detailed description can be found elsewhere^20^. Briefly, the SomaScan assay utilises a library of short single-stranded DNA molecules, which are chemically modified to specifically bind to protein targets and the relative amount of aptamers binding to protein targets is determined using DNA microarrays. To account for variation in hybridization within runs, hybridization control probes are used to generate a hybridization scale factor for each sample. To control for total signal differences between samples due to variation in overall protein concentration or technical factors such as reagent concentration, pipetting or assay timing, a ratio between each aptamer’s measured value and a reference value is computed, and the median of these ratios is computed for each of the three dilution sets (40%, 1% and 0.005%) and applied to each dilution set. Samples were removed if they were deemed by SomaLogic to have failed or did not meet our acceptance criteria of 0.25-4 for all scaling factors. In addition to passing SomaLogic QC, only human protein targets were taken forward for subsequent analysis (4,979 out of the 5284 aptamers). Aptamers’ target annotation and mapping to UniProt accession numbers as well as Entrez gene identifiers were provided by SomaLogic.

We estimated a limit of detection (LOD) for each SOMAmer reagent using a “robust estimate” method suggested by SomaLogic, based on the median plus 4.9 * median absolute deviation (MAD) signal of the blank (buffer) samples. We further defined outliers for SOMAmer and Olink measurements as being outside the median ± 5*MAD based on test sample signals and used the fraction of outliers as a variable to explain variation.

Plasma samples for a subset of 500 Fenland participants were additionally measured using 12 Olink 92-protein panels using proximity extension assays^16^. Of the 1104 Olink proteins, 1069 were unique (n=35 on >1 panel, average correlation coefficient 0.90). We imputed values below the detection limit of the assay using raw fluorescence values. Protein levels were normalized (‘NPX’) and subsequently log2-transformed for statistical analysis. A total of 15 samples were excluded based on quality thresholds recommended by Olink, leaving 485 samples for analysis.

### Protein target mapping

We mapped each candidate protein to its UniProt-ID^30^ and used those to select mapping SOMAmer reagents and Olink measures based on annotation files provided by the vendors. We further queried the UniProt database to obtain protein domain information and other characteristics of overlapping protein targets.

### Statistical analysis

We used rank-based inverse normal transformations to make protein measurements between both technologies comparable and reported Spearman rank-based and Pearson correlation coefficients as a measure of concordance between platforms.

To derive factors explaining the Spearman correlation gradient across protein targets, we created a matrix with meta-information for each protein target, including information about technical characteristics of each platform as well as characteristics of the protein target (**Fig. 2**) and used those as input for a Random-forest based feature selection approach, called Boruta-feature selection^31^. Briefly, this method employs multiple rounds of Random-forest generation and includes so-called shadow variables, which are permuted versions of the original input variables, to derive test statistics for the variable importance measure.

### Genotyping and imputation

Fenland participants were genotyped using one of three genotyping arrays: the Affymetrix UK Biobank Axiom array (OMICs, N=8994), Illumina Infinium Core Exome 24v1 (Core-Exome, N=1060) and Affymetrix SNP5.0 (GWAS, N=1402). Samples were excluded for the following reasons: 1) failed channel contrast (DishQC <0.82); 2) low call rate (<95%); 3) gender mismatch between reported and genetic sex; 4) heterozygosity outlier; 5) unusually high number of singleton genotypes or 6) impossible identity-by-descent values. Single nucleotide polymorphisms (SNPs) were removed if: 1) call rate < 95%; 2) clusters failed Affymetrix SNPolisher standard tests and thresholds; 3) MAF was significantly affected by plate; 4) SNP was a duplicate based on chromosome, position and alleles (selecting the best probeset according to Affymetrix SNPolisher); 5) Hardy-Weinberg equilibrium p<10^−6^; 6) did not match the reference or 7) MAF=0.

Autosomes for the OMICS and GWAS subsets were imputed to the HRC (r1) panel using IMPUTE4^32^, and the Core-Exome subset and the X-chromosome (for all subsets) were imputed to HRC.r1.1 using the Sanger imputation server^33^. All three arrays subsets were also imputed to the UK10K+1000Gphase3^34^ panel using the Sanger imputation server in order to obtain additional variants that do not exist in the HRC reference panel. Variants with MAF < 0.001, imputation quality (info) < 0.4 or Hardy Weinberg Equilibrium p < 10^−7^ in any of the genotyping subsets were excluded from further analyses.

### GWAS and meta-analysis

After excluding ancestry outliers and related individuals, 10,708 Fenland participants had both phenotypes and genetic data for the GWAS (OMICS=8,350, Core-Exome=1,026, GWAS=1,332). Within each genotyping subset, aptamer abundances were transformed to follow a normal distribution using the rank-based inverse normal transformation. Transformed aptamer abundances were then adjusted for age, sex, sample collection site and 10 principal components in STATA v14 and the residuals used as input for the genetic association analyses. Test site was omitted for protein abundances measured by Olink as those were all selected from the same test site. Genome-wide association was performed under an additive model using BGENIE (v1.3)^32^. Results for the three genotyping arrays were combined in a fixed-effects meta-analysis in METAL^35^. Following the meta-analysis, 17,652,797 genetic variants also present in the largest subset of the Fenland data (Fenland-OMICS) were taken forward for further analysis.

For each protein target, we used a genome-wide significance threshold of 1.004×10^−11^ (SomaScan) or 4.5 ×10^−11^ (Olink) and defined non-overlapping regions by merging overlapping or adjoining 1Mb intervals around all genome-wide significant variants (500kb either side), treating the extended MHC region (chr6:25.5–34.0Mb) as one region. We classified pQTLs as *cis*-acting instruments if the variant was less than 500kb away from the gene body of the protein encoding gene.

We performed conditional analysis as implemented in the GCTA software using the *slct* option for each genomic region - aptamer pair identified. We used a collinear cut-off of 0.1 and a p-value below 5×10^−8^ to identify secondary signals in each region. As a quality control step, we fitted a final model including all identified variants for a given genomic region using individual level data in the largest available data set (‘Fenland-OMICs’) and discarded all variants no longer meeting genome-wide significance.

To facilitate comparison between SomaScan and Olink, we repeated genetic variant – protein target associations within the same sample for which Olink was available. To account for differing sample sizes between the SomaScan data in Fenland and the varying sample sizes within SCALLOP, we recomputed p-values holding the beta estimates constant and re-estimated standard errors using the respective sample size. We considered a predicted p-value threshold of 10^−5^ to include pQTLs for consistency assessment in case there was evidence for a genome-wide signal from either approach.

### Annotation of pQTLs

For each identified pQTL we first obtained all SNPs in at least moderate LD (R^2^>0.1) using PLINK (version 2.0) and queried comprehensive annotations using the variant effect predictor software^36^ (version 98.3) using the *pick* option. For each *cis*-pQTL we checked whether either the variant itself or a proxy in the encoding gene (R^2^>0.1) is predicted to induce a change in the amino acid sequence of the associated protein, so-called protein altering variants (PAVs).

### Phenome-wide association analyses

To enable linkage to reported GWAS-variants we downloaded all SNPs reported in the GWAS catalog^37^ (19/12/2019) and pruned the list of variant-outcome associations manually to omit previous protein-wide GWASs. For each SNP identified in the present study we tested whether the variant or a proxy in LD (R^2^>0.8) has been reported to be associated with other outcomes previously.

We used the Open GWAS database^38^ to query for each genomic region association with non-proteomic phenotypes and tested for a shared genetic signal between a protein target and a phenotype with at least suggestive evidence (p<10^−6^) using statistical colocalisation^39^. We considered a posterior probability of 80% as highly likely. We repeated this analysis for all *cis*-regions from the SomaScan-based discovery with evidence for a secondary signal (p<5×10^−8^) by creating conditional summary statistics using the lead signal in the locus as additional covariate. We computed conditional association statistics using the *cond* option from GCTA-cojo to align with the identification of secondary signals.

### Expression quantitative trait loci

We obtained lead eQTLs from the most recent release of the GTEx project v8^40^ across all 49 tissues and mapped *cis*-pQTLs to *cis*-eQTLs by LD (R^2^>0.8) restricting to the respective protein-encoding gene. We further generated a simple LD-based mapping (R^2^>0.8) considering any overlap between lead pQTLs and eQTLs to allow for incorporation of *trans*-pQTLs.

### Analysis of genetic associations

We used logistic regression models to test whether variant or protein characteristics as well as technical factors were associated with the likelihood of a shared genetic region. We stratified these analyses by having a common set of shared control regions but three different sets of platform-specific regions, including regions with evidence for distinct signals within the same region (±500kB) or regions only seen when using either of the two assay platforms. We derived robust standard errors using the sandwich method. We applied log-transformation (‘apparent Kd’) or square root-transformation (number of colocalising traits, absolute effect estimate, and predicted explained variance) to reduce the impact of highly skewed factors.

To decompose the variance of measurement differences we computed the differences in rank-transformed measurements between SomaScan and Olink for each overlapping protein target. We used this variable as outcome for a variance decomposition model as implemented in R package *variancePartition* using a corresponding pQTL, age, sex, body mass index, plasma alanine aminotransferase, and estimated glomerular filtration rate as explanatory variables. We selected the only one pQTL for each overlapping pair based on a simple linear regression model explaining the differences in measurements.

Finally, we used a linear regression model to test whether the association between the Olink measure (outcome) and the SomaScan measure (exposure) differed by genotype of associated pQTLs. The resulting p-value for the interaction term between the SomaScan variable and the pQTL can be interpreted as a test of differential correlation coefficients based on genotype. We accounted for multiple testing by adopting a false discovery rate of 20%. We took a permissive approach given the small sample size (N=485) and the generally low statistical power to detect interaction terms.

We used R version 3.6.0 (R Foundation for statistical computing, Vienna, Austria) and BioRender.com for visualization of results.

## DATA AVAILABILITY

Information about the Fenland cohort is available at the study website (https://www.mrc-epid.cam.ac.uk/research/studies/fenland/information-for-researchers/), which includes a link to the MRC Epidemiology Unit metadata access portal (https://epi-meta.mrc-epid.cam.ac.uk/) which describes how data can be accessed by bona fide researchers for specified scientific purposes. Data will either be shared through an institutional data sharing agreement or arrangements will be made for analyses to be conducted remotely without the necessity for data transfer. Publicly available summary statistics for look-up and colocalisation of pQTLs were obtained from https://gwas.mrcieu.ac.uk/ and https://www.ebi.ac.uk/gwas/. We obtained genome-wide summary statistics for 90 protein targets from Folkersen et al.^8^, which are also available from the GWAS catalog (GCST90011994-GCST90012083).

## ACKNOWLEDGEMENT

The Fenland Study (10.22025/2017.10.101.00001) is funded by the Medical Research Council (MC_UU_12015/1). We are grateful to all the volunteers and to the General Practitioners and practice staff for assistance with recruitment. We thank the Fenland Study Investigators, Fenland Study Co-ordination team and the Epidemiology Field, Data and Laboratory teams. We further acknowledge support for genomics from the Medical Research Council (MC_PC_13046). Proteomic measurements were supported and governed by a collaboration agreement between the University of Cambridge and Somalogic. JCZ is supported by a 4-year Wellcome Trust PhD Studentship and the Cambridge Trust, CL, EW, and NJW are funded by the Medical Research Council (MC_UU_12015/1). NJW is a NIHR Senior Investigator. ADH is an NIHR Senior Investigator and supported by the UCL Hospitals NIHR Biomedical Research Centre and the UCL BHF Research Accelerator (AA/18/6/34223). We thank Philippa Pettingill, Ida Grundberg, Klev Diamanti, and Andrea Ballagi for advice and comments on an earlier draft of this manuscript.

## AUTHOR CONTRIBUTIONS

MP and CL designed the analysis and drafted the manuscript. MP, EW, and JL analysed the data. JCZ and EO provided bioinformatic characterization of protein targets and mapped pQTLs to eQTLs. NK and EO performed quality control of proteomic measurements. ADH and SAW provided critical review and intellectual contribution to the discussion of results. NJW is PI of the Fenland cohort. All authors contributed to the interpretation of the results and critically reviewed the manuscript.

