## Supplementary Figures 1-4 for "Cross-platform proteomics to advance genetic prioritisation strategies"

^5^Health Data Research UK

^6^SomaLogic, Inc., Boulder, CO, USA

**FIGURES**

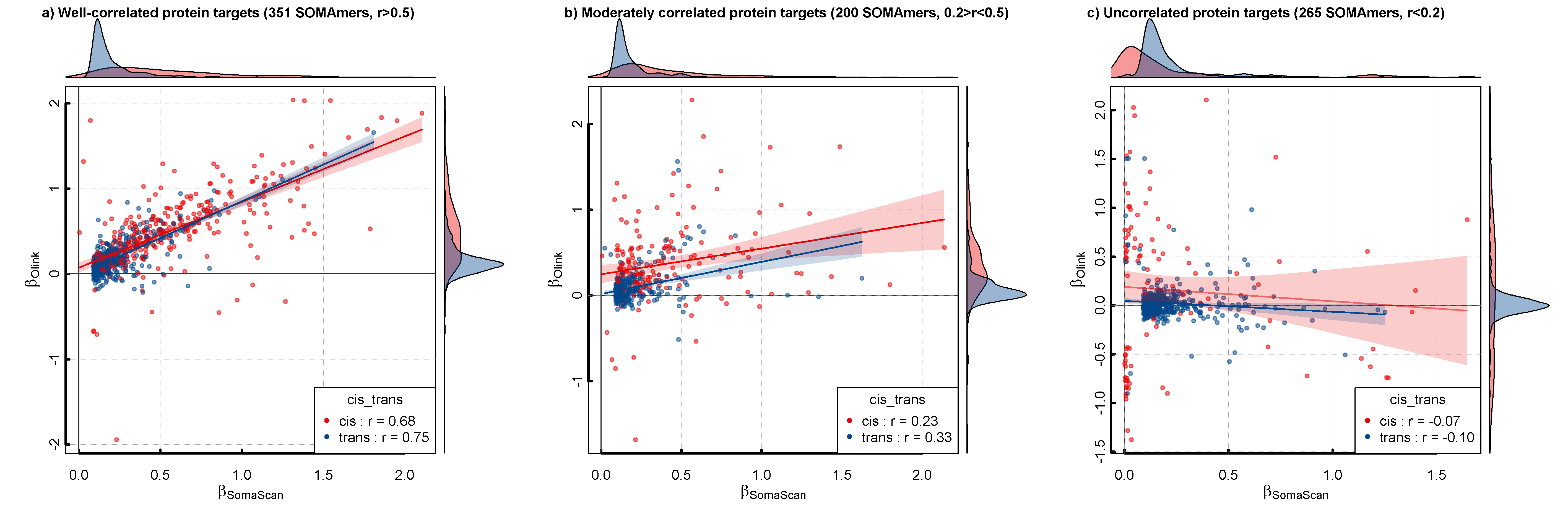

**Supplementary Figure 1** Stratification of effect estimate correlations for genetic variants associated with either the SomaScan-based or Olink-based discovery. Colouring is based on the genomic location of genetic variants. Red indicates variants close to the protein encoding gene (cis, ±500kb) and blue otherwise.

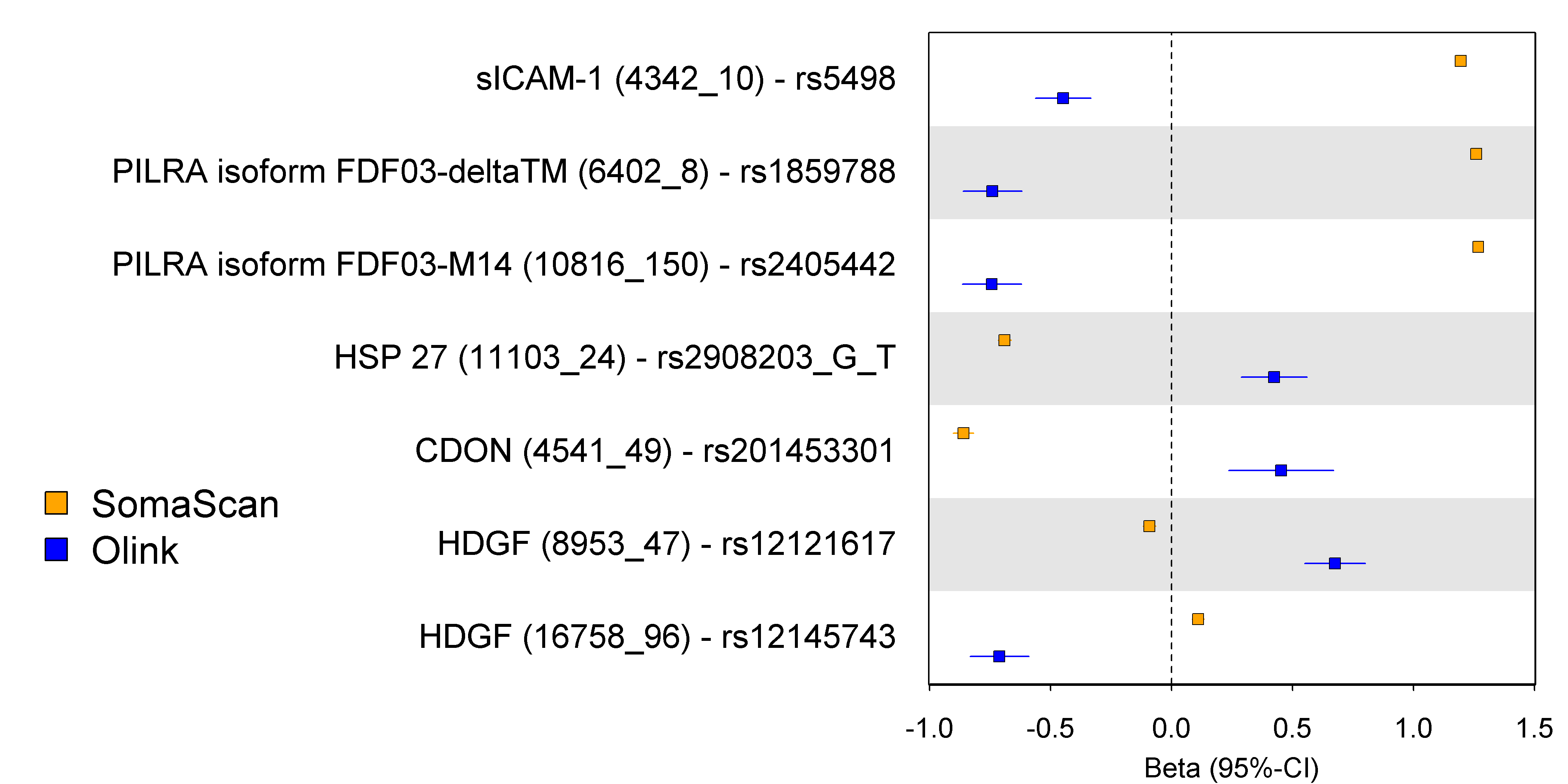

**Supplementary Figure 2** Effect estimates for protein quantitative trait loci common to the SomaScan assay and Olink for protein targets with discordant effect estimates. Numbers in brackets are identifiers for SOMAmer reagents. Each row contains the effect of the genetic variant for the SOMAmer reagent (orange) and the corresponding Olink measurement (blue).

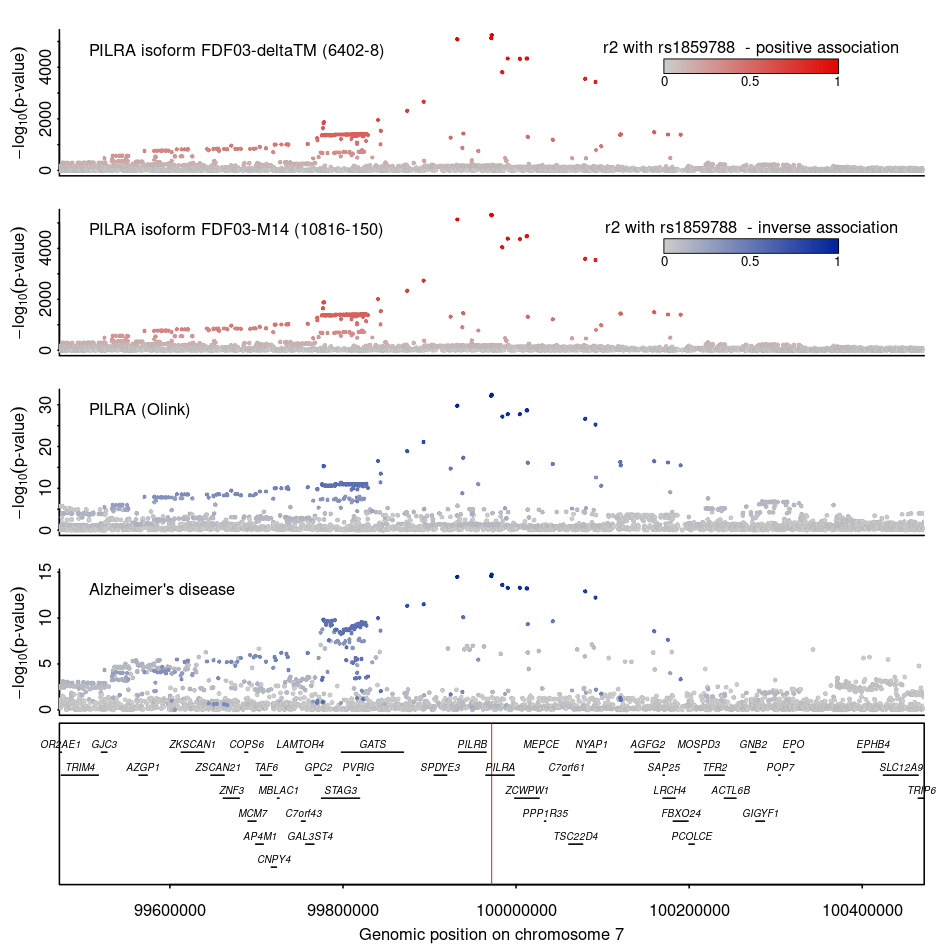

**Supplementary Figure 3** Regional association plots for Paired immunoglobulin-like type 2 receptor alpha measured by SomaScan (top rows) and Olink, as well as for Alzheimers disease centred around a coloclasing signal for the missense variant rs1859788 wihtin *PILRA* (p.G78R). Colours indicate direction of effect for the A-allele of rs1859788 on the respective trait (blue – inverse, red - positive) and shading indicates linkage disequilibrium (R^2^) with the lead variant at the locus. The red line in the gene panel indicates the position of the variant. P-values for protein measures were derived from genome-wide association analysis from the Fenland cohort as described in the main text, whereas summary statistics for Alzheimer’s disease were obtained from Jansen et al. 2019^1^.

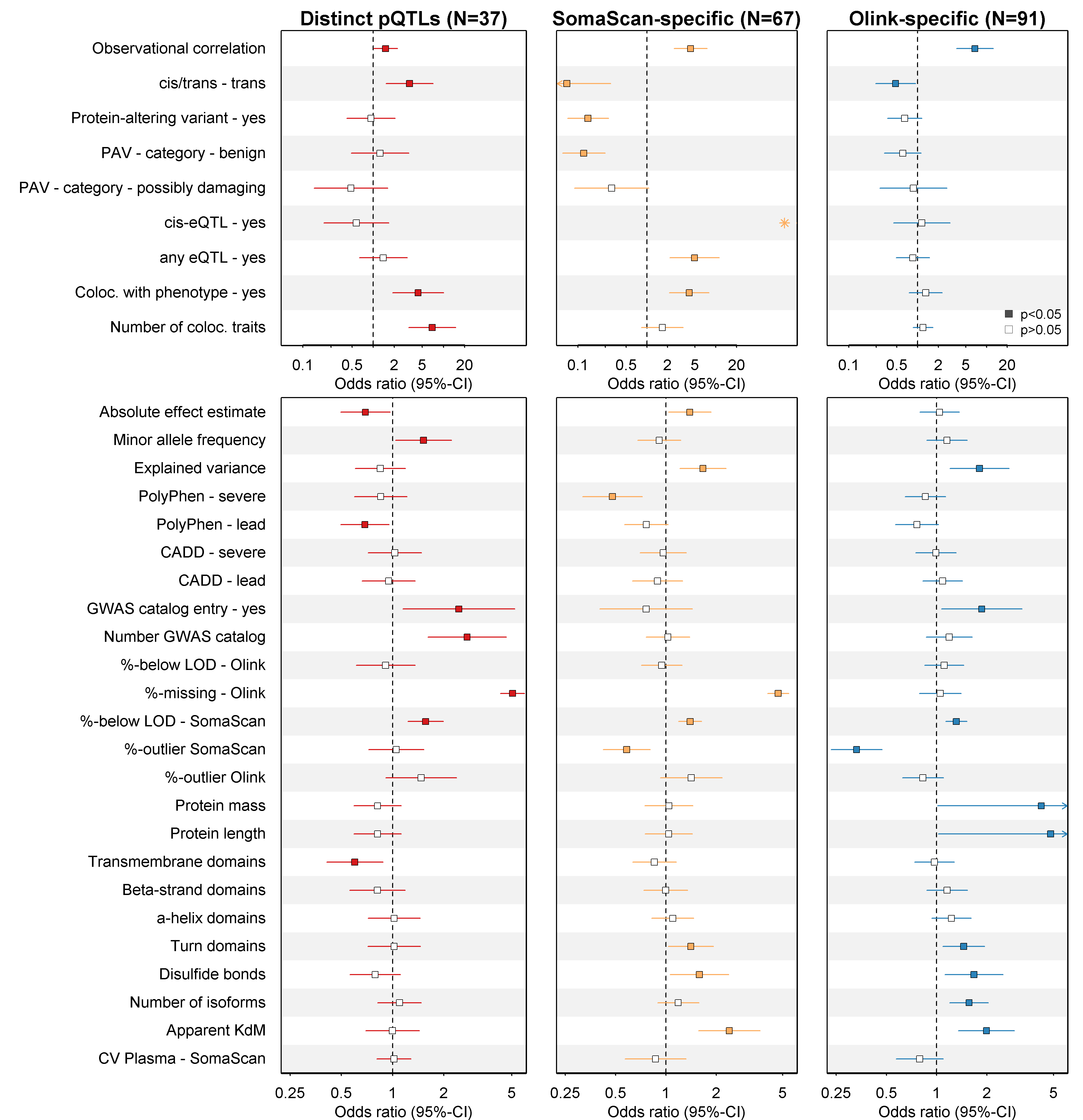

**Supplementary Figure 4** **Factors associated with pQTLs consistent across platforms compared to three sets controls**. Odds ratios and 95%-confidence intervals for factors associated with cross-platform protein quantitative trait loci (pQTL) across the SomaScan v4 and Olink assays. The panels 318 variant – protein target pairs (120 shared, 198 platform-specific) from the SCALLOP CVD-I effort. Colours indicate the reason for inconsistency and filled rectangles indicate significant p-values (p<0.05). The blue star indicates non-convergence of the logistic regression model due to perfect separation, that is, none of the pQTLs unique to SomaScan had a *cis*-eQTL, whereas eleven among the consistent set did. PAV = protein altering variant; eQTL = expression quantitative trait loci; Coloc. = colocalisation; GWAS = genome-wide association analysis; LOD = limit of detection; KdM = estimated apparent dissociation constant (Kd) of SOMAmer reagents in molar units (M)
